# Using Adaptive Psychophysics to Identify the Neural Network Reset Time in Sub-second Interval Timing

**DOI:** 10.1101/2021.02.08.430050

**Authors:** Renata Sadibolova, Stella Sun, Devin B. Terhune

**Affiliations:** Department of Psychology, Goldsmiths, University of London

**Keywords:** state-dependent network, breakpoint, time perception, temporal discrimination, adaptive psychophysics

## Abstract

State dependent network models of sub-second interval timing propose that duration is encoded in states of neuronal populations that need to reset prior to a novel timing operation in order to maintain optimal timing performance. Previous research has shown that the approximate boundary of this reset interval can be inferred by varying the interstimulus interval between two to-be-timed intervals. However, the estimated boundary of this reset interval is broad (250-500ms) and remains underspecified with implications for the characteristics of state dependent network dynamics subserving interval timing. Here we probed the interval specificity of this reset boundary by manipulating the interstimulus interval between standard and comparison intervals in two sub-second auditory duration discrimination tasks (100 and 200ms) and a control (pitch) discrimination task using adaptive psychophysics. We found that discrimination thresholds improved with the introduction of a 333ms interstimulus interval relative to a 250ms interstimulus interval in both duration discrimination tasks, but not in the control task. This effect corroborates previous findings of a breakpoint in the discrimination performance for sub-second stimulus interval pairs as a function of an incremental interstimulus delay but more precisely localizes the minimal interstimulus delay range. These results suggest that state dependent networks subserving sub-second timing require approximately 250-333ms for the network to reset in order to maintain optimal interval timing.

**New & Noteworthy:** The state-dependent-network model considers interval timing as an intrinsic ability of neuronal populations to track the temporal evolution of their collective state. However, the time-dependent nature of neuronal properties imposes constraints on a maximum encodable interval and on the processing of intervals that are presented before the network resets to its baseline state. Investigating temporal discrimination thresholds as a function of variable inter-stimulus-intervals, we showed that the network reset time is between 250 and 333ms.

## Introduction

The *state-dependent network* (SDN) model (Buonomano & Merzenich, 1998; Karmarkar & Buonomano, 2007; Paton & Buonomano, 2018) proposes that interval timing on a scale of milliseconds to seconds is encoded in states of neuronal populations analogous to the evolving state of a liquid surface which has been disturbed by throwing in an object (Buonomano et al., 2009). One consequence of this model is that a network needs to dynamically reset in order to facilitate optimal timing: prior to network resetting, a timing operation will be deleteriously affected just as throwing in a second object before the liquid returns to its baseline state creates a distorted spatiotemporal pattern of the ripples on its surface. Previous studies suggest that neural networks supporting interval discrimination require between 250-500ms to return to their initial state (Buonomano et al., 2009; Karmarkar & Buonomano, 2007). However, it remains to be investigated which interval within that range corresponds with the network reset time and thus after how long the first stimulus ceases to influence the timing of subsequent stimuli.

The SDN model is a member of a class of computational models (Buonomano & Maass, 2009; Buonomano & Merzenich, 1995; Cueva et al., 2020; Laje & Buonomano, 2013) put forward to explain how neural networks may encode temporal stimuli without an explicit linear measure of duration such as the pulses of an oscillator (Allman et al., 2014; Matell & Meck, 2004) or the ramping firing rate of neurons (Durstewitz, 2003; Simen et al., 2011). Rather, SDN models propose that populations of neurons time intervals intrinsically by tracking the temporal evolution of the collective state of the network (Karmarkar & Buonomano, 2007). According to these models, neural networks embody an interplay of responding neurons (*active state*) constrained by their time-dependent cellular and synaptic properties with known constants in tens to hundreds of milliseconds (*hidden state*) (Buonomano & Maass, 2009; Paton & Buonomano, 2018). By tracking the state-space trajectories of a computational neural network, Karmarkar and Buonomano (2007) found that it had returned to the neighborhood of its initial state within 750ms from the offset of a stimulus interval. Whether the new trajectories for subsequent stimulus intervals reproduced the observed trajectories was contingent on the reset to the baseline state. When it was not permitted due to a rapid presentation of another stimulus interval within 250ms from the offset of the previous stimulus, an altered state-space trajectory, and correspondingly diminished temporal performance, were observed.

In keeping with the premise that both the new stimulus interval and the ongoing network state (i.e., the context imposed by the previous interval) determine the response on a given trial, several psychophysical studies sought to assess the impact of the preceding distractor intervals on temporal performance (Buonomano et al., 2009; Karmarkar & Buonomano, 2007; Spencer et al., 2009). For example, Karmarkar and Buonomano (2007) developed a “reset task” consisting of interleaved trials with a single target interval bound by two tones and a distractor and target interval-pair demarcated by three tones. The target intervals of 100ms, unlike those of 1000ms, were characterized by poorer discrimination when preceded by distractor intervals. Additionally, in another study (Spencer et al., 2009), the detrimental effect of a distractor on temporal discrimination was replicated for a 100ms stimulus interval but was not observed when either the target or the *distractor* stimulus interval increased to 300ms. One interpretation of these observations is that the network resets after a specific interval, i.e. at a maximum duration that the network might be capable of representing (Spencer et al., 2009).

It has been hypothesized that the boundary beyond which sub-second interval timing no longer relies on state-dependent computations can be identified as the inter-stimulus-interval (ISI) between the target stimulus interval pair that is associated with an improvement in the temporal discrimination threshold (Buonomano et al., 2009). In order to evaluate this, Buonomano and colleagues (2009) had participants discriminate two tone intervals (standard and comparison intervals), in blocks of trials defined by different ISIs (50, 250, 500, 750, and 1000ms). The boundary for the putative reset interval was observed between 250-500ms, as reflected by superior duration discrimination thresholds in longer ISI conditions. Nevertheless, the interval of this boundary remains underspecified.

The aim of this study was to build upon previous research (Buonomano et al., 2009; Karmarkar & Buonomano, 2007) and more precisely identify the interval boundary of network resetting in the range of 250 to 500ms. Toward this end, we measured duration discrimination thresholds for 100ms and 200ms standard stimulus intervals, and pitch discrimination threshold as a control task, in conditions with different ISIs (range: 250 to 583ms). We expected a changepoint in duration discrimination thresholds across ascending ISIs that would generalize across the two interval conditions. This changepoint was hypothesized to reflect the boundary of the network reset for interval timing and thus was not expected in the control task.

## Method

### Participants

Forty right-handed (Oldfield, 1971) individuals participated in this study and 38 participants were included in the analyses after removing two multivariate outliers (82% female, 18% male, age range: 20-34; *M*=25.63, *SD*=3.66; years of higher education range: 0-8, *M*=3.76; *SD=2.02* [four missing]). The sample size was determined *a priori* using G-power (v. 3.1.9.3; Faul et al., 2009) with a repeated-measures analysis of variance and parameters α=.05, 1-β=80%, and η_p_^2^=.08 (Buonomano et al., 2009), yielding a sample size of *N*=36. In order to account for attrition, we intended to include 40 participants and increased this to 41 when one participant, whose data was subsequently excluded, was unable to understand the tasks. All procedures were approved by the departmental Ethics committee at Goldsmiths, University of London.

### Materials

#### Duration and pitch discrimination tasks

Participants completed two duration discrimination tasks and a pitch discrimination task for the purpose of investigating discrimination thresholds as a function of ISI (250, 333, 417, 500 and 583ms). In all tasks, the trial sequence consisted of a pre-stimulus interval (500ms), a pair of tones separated by an ISI, a post-stimulus interval (500ms), and a visual response prompt (Figure 1). The two tones consisted of a fixed standard tone and a comparison tone that varied adaptively with performance, with order of presentation counterbalanced within blocks. At the prompt, participants judged whether the second tone was shorter or longer than the first tone (S L or L S [S=shorter; L=longer]) or of a lower or higher pitch (L H or H L [L=lower; H=higher]). Participants responded with their right index and middle fingers on the left and right arrow keys of a keyboard, respectively, with the response-key mappings (S L vs. L S and L H vs. H L) counterbalanced across participants. Auditory stimuli were generated in MATLAB 2018b (MathWorks, Natick) in real-time using the PsychPortAudio function of Psychtoolbox-3 (Brainard, 1997; Kleiner et al., 2007), using the Windows 7 WASAPI sound device. We sampled stereo sounds at 48 kHz default rate and presented them via the headphones at a constant intensity level (set to 0.01 programmatically and 70% in Windows sound settings).

**Figure 1:**
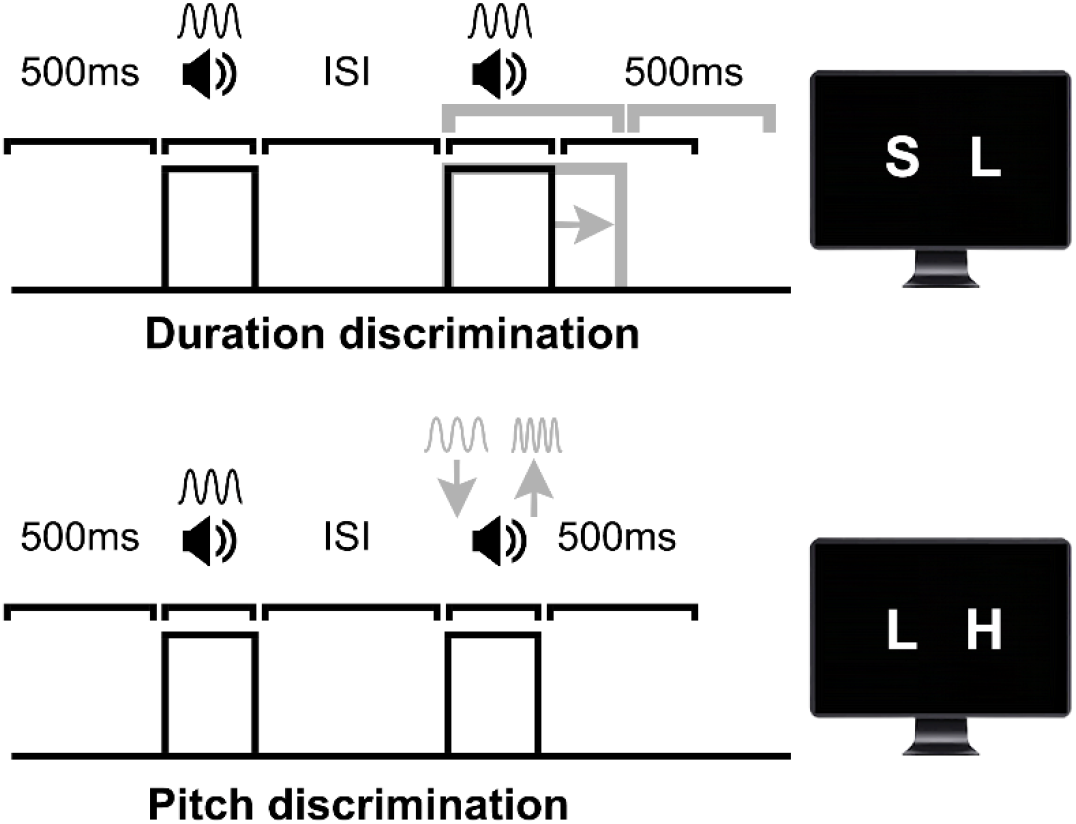
Diagrams of experimental tasks. All trials comprised a pre-stimulus interval (500ms), a pair of tones separated by an ISI (250, 333, 417, 500 or 583ms, varied at block level), a fixed post-stimulus interval (500ms), and the response prompt. Participants estimated in three two-alternative forced-choice (2AFC) tasks if the second stimulus was shorter or longer (duration discrimination tasks) or lower or higher in pitch (pitch discrimination task) compared to the first stimulus. The standard stimulus (the first of two tones in the diagrams) was fixed in each task: 100ms, 200ms, and 1kHz. The duration of the comparison stimulus (duration discrimination) and pitch of the comparison stimulus (pitch discrimination) were adaptively adjusted based on the performance on a trial-by-trial basis (grey arrows and lines). The standard-stimulus presentation order in the experiment was randomized within blocks.

In the two duration discrimination tasks, the standard stimulus duration (*d*) was 100ms and 200ms, respectively, whereas the comparison stimulus lasted *d*+Δ*d*. In these two tasks, pitch remained constant (1kHz). Analogously in the pitch discrimination task, the pitch for standard and comparison stimuli was respectively *p*=1kHz and *p*+Δ*p*, and stimulus intervals were fixed at 100ms. The change (Δ) in duration or pitch was always a positive value and it was computed on trial-by-trial basis as a psychometric threshold using a Bayesian adaptive staircase method (Ψ-marginal algorithm) implemented in the Palamedes toolbox for MATLAB (Kingdom & Prins, 2016). This method optimizes both the sampling and estimation of the target psychometric parameter(s), including participant’s responses into a prior distribution that affects subsequent values tested.

Our objective was to identify the Δ for a reliably discriminable stimulus-pair (threshold, alpha parameter), with a subsidiary assessment of temporal precision (beta) and attentional lapse rate (lambda). The guess rate (gamma) was fixed at 50%. The psychometric function parameters were updated after each trial response (correct vs. incorrect) and responses were fitted with a logistic Weibull function. The dependent measure was each block’s final discrimination threshold, i.e., the comparison stimulus Δ with 75% probability of a correct response. We constrained the prior beta and lambda parameters to values from zero to four in steps of .1 and zero to .2 in steps of .02, respectively. The initial prior alpha range was 100 to 300ms and 200 to 400ms (both in 1ms steps) respectively for the 100ms and 200ms standards and 1kHz to 1.5kHz (5Hz steps) for the 1kHz standard. These upper boundaries were extended for acceptable comparison stimulus range by 300ms in duration discrimination tasks and 2kHz for pitch discrimination.

### Procedure

Following general instructions, the experimenter confirmed with each participant that the stimulus volume was well above the detection threshold yet within a safe audible range. Prior to each task, participants completed ten practice trials with randomly selected standard-comparison stimulus pairs. Participants subsequently completed five consecutive blocks for each of the three tasks in randomized order (each corresponding to a unique ISI: 250, 333, 417, 500 and 583ms) resulting in 15 blocks. Task order was counterbalanced across participants. To avoid fatigue, short breaks after each block were encouraged. The entire experiment took approximately 60 minutes.

### Analyses

Two participants were removed as multivariate outliers with Mahalanobis distance values > 31.02, *p*=.001. Data were reliably characterized by a departure from normality (Shapiro-Wilk test *p*<.05, Figure 2D) and thus they were analyzed using nonparametric Friedman tests and Wilcoxon signed-rank tests (in IBM SPSS Statistics software v.24). P-values for the latter tests were adjusted using a Holm-Bonferroni multiple-comparison correction (Holm, 1979). We report Kendal W (*r*_τ_) effect size for the Friedman tests and *r* (*r* = *z*/√*N*; (Pallant, 2007)) for the Wilcoxon tests. Bayes factors were not computed due to violations of normality (Dienes, 2014; Rouder et al., 2012; Wetzels et al., 2012).

**Figure 2:**
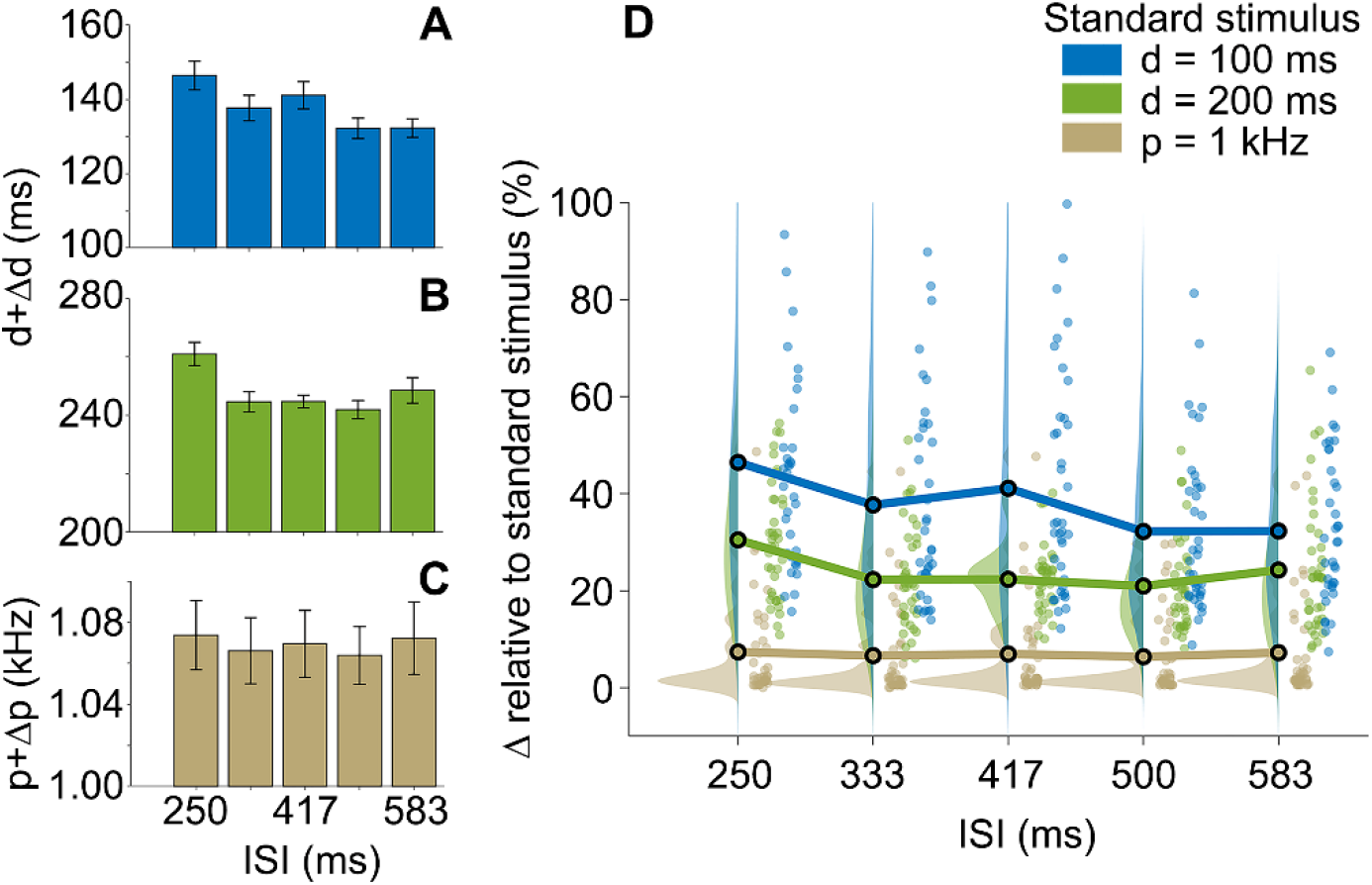
Duration (d) discrimination and pitch (p) discrimination as a function of the ISI (ms) between standard and comparison stimuli. **(A-C)** Δ (75% discrimination threshold) for different ISIs in duration and pitch discrimination tasks. Error bars indicate standard error of the mean (SEM). **(D)** Δ scaled by the respective standard stimulus. Marginal plots show the kernel density distributions and individual participant data in each condition (Allen et al., 2019).

## Results

Previous research suggests that the duration discrimination varies according to the ISI between two intervals, such that a rapid succession (short ISI) is associated with poorer performance (higher discrimination threshold) (Buonomano et al., 2009). As can been seen in Figure 2A-B, performance patterns on the temporal discrimination tasks generally conform to this pattern as the highest discrimination thresholds (*d+Δd*) in both duration discrimination tasks were observed for the shortest ISI (250ms). These effects were reflected in significant main effects of ISI on duration discrimination thresholds in both tasks with similar effect sizes, 100ms: *χ^2^_F_*(4)=25.50, *p*<.001, *r_τ_*=.17, and 200ms: *χ^2^_F_*(4)=20.78, *p*<.001, *r_τ_*=.14. As anticipated, a corresponding main effect of ISI was not observed for pitch discrimination thresholds, *χ^2^_F_*(4)=3.87, *p*=.42, *r_τ_*=.03, showing relative uniformity across ISIs (Figure 2C-D). These results suggest that duration discrimination thresholds selectively vary as a function of ISI.

In order to identify the ISI at which the earlier stimulus ceases to interfere, we conducted four planned comparisons of duration discrimination thresholds between the adjacent ISIs in each duration discrimination task. In the 100ms standard task, duration discrimination thresholds were higher in the 250ms (*Mdn*=143.60ms) than in the 333ms (*Mdn*=128.40) ISI condition, although this difference was only observed at a trend-level, *Z*=−2.47, *p*=.08, *r*=−.40 (Figure 2A). There was no significant difference between thresholds in the 333ms and 417ms ISI conditions (*Mdn*=133.00), *Z*=−0.50, *p*=1.00, *r*=−.08. By contrast, discrimination thresholds were significantly lower (improved) in the 500ms (*Mdn*=126.75) relative to the 417ms ISI condition, *Z*=−3.25, *p*=.01, *r*=−.53 and were not significantly different between the 500 and 583ms (*Mdn*=130.05) ISI conditions, *Z*=−0.15, *p*=1.00, *r*=−.02. This pattern of results replicates previous observations (Buonomano et al., 2009) and suggests that the network requires less than 500ms to reset. However, there was some ambiguity regarding the precise window of this reset with weak evidence for an early reset (250-333ms) and additional evidence for a later reset (417-500ms). Such ambiguity was not present in the 200ms standard task where there was clear evidence for an earlier boundary in alignment with the former effect. In particular, duration discrimination thresholds were significantly greater in the 250ms (*Mdn*=259.80ms) than in the 333ms (*Mdn*=242.45ms) ISI condition, *Z*=−3.48, *p*=.01, *r*=−.56. Duration discrimination thresholds remained relatively stable across the remaining ISI conditions (Figure 2B), 333ms vs. 417ms (*Mdn*=244.75ms), *Z*=−0.66, *p*=1.00, *r*=−.11, 417ms vs. 500ms (*Mdn*=237.90ms), *Z*=−1.29, *p*=1.00, *r*=−.21, and 500ms vs. 583ms (*Mdn*=245.50ms), *Z*=−1.31, *p*=1.00, *r*=−.21. Cumulatively, these results suggest a boundary for this network reset between 250 and 333ms.

Additional analyses of discrimination thresholds with Δ scaled by the standard stimulus (Figure 2D) corroborated that the temporal discrimination task was more difficult in the 100ms standard stimulus condition (*Mdn*=34.85%) compared to the 200ms standard stimulus condition (*Mdn*=23.97%), *Z*=−5.36, *p*<.001, *r*=−.87. In order to explore the interaction of the ISI and standard stimulus conditions, particularly the later reset for the 100ms vs. the 200ms standard condition, we subtracted the scaled Δ in the 100ms condition from those in the 200ms condition. A Friedman test on the difference in scaled Δ did not yield a significant interaction effect, *X^2^_F_*(4)=6.80, *p*=.15, *r_τ_*=.05. These results suggest that the two standard stimulus conditions did not significantly differ in the reset interval.

## Discussion

SDN models of interval timing propose that optimal sub-second interval timing requires resetting of neuronal networks encoding stimulus duration prior to a new timing operation (Karmarkar & Buonomano, 2007; Paton & Buonomano, 2018; Spencer et al., 2009). This study sought to estimate the putative network reset interval by varying the ISI between comparison and standard intervals in a sub-second auditory interval discrimination task (Buonomano et al., 2009). We found that interval discrimination thresholds significantly improved (decreased) in the 200ms standard condition and suggestively improved in the 100ms standard condition when the ISI increased from 250 to 333ms, with moderate effect sizes in both cases. By contrast, the analogous comparisons for pitch discrimination thresholds yielded non-significant results. These findings are consistent with previous research (Buonomano et al., 2009; Karmarkar & Buonomano, 2007) suggesting that the network reset interval is between 250 and 500ms, warranting further research on the characteristics and dynamics of network resetting in sub-second interval timing.

Computational studies of SDNs have conventionally included a constraint that the physiological mechanisms are of limited temporal extent after which the network resets (Buonomano & Merzenich, 1998; Karmarkar & Buonomano, 2007). Germane behavioral evidence suggests that SDNs subserving interval timing reset between 250 and 500ms (Buonomano et al., 2009). This inference was made on the basis of a decrease in discrimination thresholds for pairs of intervals separated by 500ms, relative to 250ms ISIs. Our results corroborate this time window for the putative SDN reset (250-500ms) and show that this effect generalizes to a 200ms standard interval stimulus condition. Moreover, we further expanded upon the previous findings through the inclusion of a greater number of ISIs during the putative breakpoint window (250, 333, 417, 500ms) in order to permit greater precision in the estimate of the network reset interval. As a result, we identified a narrower reset time window of 250-333ms which aligns with the observation of improved temporal performance in a ‘reset task’ when either the distractor interval or immediately following *target* interval increased from 100 to 300ms (Spencer et al., 2009).

Although the reset time, and therefore the inability to accommodate longer intervals, have previously been considered to be a limitation of the applicability of the SDN model in timing (Spencer et al., 2009), the presence of a mechanism dedicated to the processing of sub-second intervals is congruent with more recent advances in the interval timing literature. For instance, Rammsayer and Pichelmann (2018) introduced a conceptual model of sub- and supra-second timing, arguing for distinct modality-specific neurocognitive mechanisms subserving the timing of brief intervals (below ~100-500ms) that gradually gives way to amodal mechanisms responsible for the processing of longer intervals. Indeed, the time-dependent changes in the state of neural networks are likely to incorporate modality-specific neural codes for brief intervals but much less so for longer intervals recruiting executive functions (Paton & Buonomano, 2018). Numerous studies, reporting psychopharmacological (Rammsayer, 1993; Rammsayer & Vogel, 1992), psychophysical (Karmarkar & Buonomano, 2007; Rammsayer et al., 2015), neuroimaging (Lewis & Miall, 2003; Wiener et al., 2010) or genetic (Wiener et al., 2011) evidence, add further weight to the argument of distinct timing mechanisms in the millisecond-to-second range. However, the boundary between these putative timing systems, and the nature of a transition between them, remains controversial. Provided that neural networks are responsible for the timing of very brief millisecond intervals, our findings would suggest that the boundary between sub-second timing systems is within the 250-333ms window.

Although the present results provide a more refined estimate of the network reset interval window than previous research, the characteristics of this reset interval require further specification. Further research is required to more precisely delineate the window of this reset interval. In particular, further research would benefit from adaptively varying the ISI between standard and comparison intervals in order to derive a more precise estimate of the network reset interval. The present work suggests that this interval will be observed between 250 and 333ms. It will also be important to determine whether this reset interval generalizes across sensory modalities or covaries with superior temporal precision in auditory relative to visual timing (Penney et al., 2000). A further outstanding question concerns the role of the network reset interval in differentiating timing mechanisms for sub-second and supra-second intervals. Our evidence suggests that the breakpoint that co-occurs with the network reset interval is specific to sub-second interval timing. Nevertheless, it remains understudied whether the shift to different mechanism for longer intervals is abrupt or gradual. Further work would therefore benefit from using the present approach to probe the division between sub-second and supra-second timing.

In conclusion, previous research suggests that interval timing engages disparate neural mechanisms, depending on the timescale and computational requirements of the task (Paton & Buonomano, 2018). The SDN model represents a category of intrinsic timing models implicated in the encoding of brief sub-second intervals by means of trajectories in neuronal space. Whereas further research is required to disentangle the contributions of different timing mechanisms, our results build upon previous research (Buonomano et al., 2009; Karmarkar & Buonomano, 2007) and provide a more precise estimate of the temporal window of the SDN reset and suggest that approximately 250-333ms is required for the network to reset in order to facilitate optimal interval timing.

## Acknowledgements

This research was partly funded by Biological and Biotechnology Sciences Research Council, grant number: BB/R01583X/1.

## References

Allen, M., Poggiali, D., Whitaker, K., Marshall, T. R., & Kievit, R. A. (2019). Raincloud plots: a multi-platform tool for robust data visualization. Wellcome Open Research, 4, 63. https://doi.org/10.12688/wellcomeopenres.15191.1

Allman, M. J., Teki, S., Griffiths, T. D., & Meck, W. H. (2014). Properties of the Internal Clock: First- and Second-Order Principles of Subjective Time. Annual Review of Psychology, 65(1), 743–771. https://doi.org/10.1146/annurev-psych-010213-115117

Brainard, D. H. (1997). The Psychophysics Toolbox. Spatial Vision, 10(4), 433–436. https://doi.org/10.1163/156856897X00357

Buonomano, D. V., Bramen, J., & Khodadadifar, M. (2009). Influence of the interstimulus interval on temporal processing and learning: testing the state-dependent network model. Philosophical Transactions of the Royal Society B: Biological Sciences, 364(1525), 1865–1873. https://doi.org/10.1098/rstb.2009.0019

Buonomano, D. V., & Maass, W. (2009). State-dependent computations: spatiotemporal processing in cortical networks. Nature Reviews Neuroscience, 10(2), 113–125. https://doi.org/10.1038/nrn2558

Buonomano, D. V., & Merzenich, M. (1995). Temporal information transformed into a spatial code by a neural network with realistic properties. Science, 267(5200), 1028–1030. https://doi.org/10.1126/science.7863330

Buonomano, D. V., & Merzenich, M. M. (1998). CORTICAL PLASTICITY: From Synapses to Maps. Annual Review of Neuroscience, 21(1), 149–186. https://doi.org/10.1146/annurev.neuro.21.1.149

Cueva, C. J., Saez, A., Marcos, E., Genovesio, A., Jazayeri, M., Romo, R., Salzman, C. D., Shadlen, M. N., & Fusi, S. (2020). Low-dimensional dynamics for working memory and time encoding. Proceedings of the National Academy of Sciences, 117(37), 23021–23032. https://doi.org/10.1073/pnas.1915984117

Dienes, Z. (2014). Using Bayes to get the most out of non-significant results. Frontiers in Psychology, 5(July), 1–17. https://doi.org/10.3389/fpsyg.2014.00781

Durstewitz, D. (2003). Self-Organizing Neural Integrator Predicts Interval Times through Climbing Activity. The Journal of Neuroscience, 23(12), 5342–5353. https://doi.org/10.1523/JNEUROSCI.23-12-05342.2003

Faul, F., Erdfelder, E., Buchner, A., & Lang, A. (2009). Statistical power analyses using G*Power 3.1: Tests for correlation and regression analyses. Behavior Research Methods, 41(4), 1149–1160. https://doi.org/10.3758/BRM.41.4.1149

Holm, S. (1979). A simple sequentially rejective multiple test procedure. Scandinavian Journal of Statistics.

Karmarkar, U. R., & Buonomano, D. V. (2007). Timing in the Absence of Clocks: Encoding Time in Neural Network States. Neuron, 53(3), 427–438. https://doi.org/10.1016/j.neuron.2007.01.006

Kingdom, F. A. A., & Prins, N. (2016). Psychophysics: A Practical Introduction (Second Edi). Elsevier. https://doi.org/10.1016/C2012-0-01278-1

Kleiner, M., Brainard, D. H., Pelli, D. G., Broussard, C., Wolf, T., & Niehorster, D. (2007). What’s new in Psychtoolbox-3? Perception.

Laje, R., & Buonomano, D. V. (2013). Robust timing and motor patterns by taming chaos in recurrent neural networks. Nature Neuroscience, 16(7), 925–933. https://doi.org/10.1038/nn.3405

Lewis, P.., & Miall, R.. (2003). Brain activation patterns during measurement of sub- and supra-second intervals. Neuropsychologia, 41(12), 1583–1592. https://doi.org/10.1016/S0028-3932(03)00118-0

Matell, M. S., & Meck, W. H. (2004). Cortico-striatal circuits and interval timing: coincidence detection of oscillatory processes. Cognitive Brain Research, 21(2), 139–170. https://doi.org/10.1016/j.cogbrainres.2004.06.012

Oldfield, R. C. (1971). The assessment and analysis of handedness: The Edinburgh inventory. Neuropsychologia, 9(1), 97–113. https://doi.org/10.1016/0028-3932(71)90067-4

Pallant, J. (2007). SPSS Survival Manual (3rd Ed.). McGraw Hill Open University Press.

Paton, J. J., & Buonomano, D. V. (2018). The Neural Basis of Timing: Distributed Mechanisms for Diverse Functions. Neuron, 98(4), 687–705. https://doi.org/10.1016/j.neuron.2018.03.045

Penney, T. B., Gibbon, J., & Meck, W. H. (2000). Differential effects of auditory and visual signals on clock speed and temporal memory. Journal of Experimental Psychology: Human Perception and Performance, 26(6), 1770–1787. https://doi.org/10.1037//0096-1523.26.6.1770

Rammsayer, T. H. (1993). On dopaminergic modulation of temporal information processing. Biological Psychology, 36(3), 209–222. https://doi.org/10.1016/0301-0511(93)90018-4

Rammsayer, T. H., Borter, N., & Troche, S. J. (2015). Visual-auditory differences in duration discrimination of intervals in the subsecond and second range. Frontiers in Psychology, 6(OCT), 1626. https://doi.org/10.3389/fpsyg.2015.01626

Rammsayer, T. H., & Pichelmann, S. (2018). Visual-auditory differences in duration discrimination depend on modality-specific, sensory-automatic temporal processing: Converging evidence for the validity of the Sensory-Automatic Timing Hypothesis. Quarterly Journal of Experimental Psychology, 71(11), 2364–2377. https://doi.org/10.1177/1747021817741611

Rammsayer, T. H., & Vogel, W. H. (1992). Pharmacologic Properties of the Internal Clock Underlying Time Perception in Humans. Neuropsychobiology, 26(1–2), 71–80. https://doi.org/10.1159/000118899

Rouder, J. N., Morey, R. D., Speckman, P. L., & Province, J. M. (2012). Default Bayes factors for ANOVA designs. Journal of Mathematical Psychology, 56(5), 356–374. https://doi.org/10.1016/j.jmp.2012.08.001

Simen, P., Balci, F., deSouza, L., Cohen, J. D., & Holmes, P. (2011). A model of interval timing by neural integration. Journal of Neuroscience, 31(25), 9238–9253. https://doi.org/10.1523/JNEUROSCI.3121-10.2011

Spencer, R. M. C., Karmarkar, U., & Ivry, R. B. (2009). Evaluating dedicated and intrinsic models of temporal encoding by varying context. Philosophical Transactions of the Royal Society B: Biological Sciences, 364(1525), 1853–1863. https://doi.org/10.1098/rstb.2009.0024

Wetzels, R., Grasman, R. P. P. P. P. P., & Wagenmakers, E.-J. (2012). A Default Bayesian Hypothesis Test for ANOVA Designs. The American Statistician, 66(2), 104–111. https://doi.org/10.1080/00031305.2012.695956

Wiener, M., Lohoff, F. W., & Coslett, H. B. (2011). Double Dissociation of Dopamine Genes and Timing in Humans. Journal of Cognitive Neuroscience, 23(10), 2811–2821. https://doi.org/10.1162/jocn.2011.21626

Wiener, M., Turkeltaub, P., & Coslett, H. B. (2010). The image of time: A voxel-wise meta-analysis. NeuroImage, 49(2), 1728–1740. https://doi.org/10.1016/j.neuroimage.2009.09.064

